# *N*6-methyladenosine dynamics during early vertebrate embryogenesis

**DOI:** 10.1101/528422

**Authors:** Håvard Aanes, Dominique Engelsen, Adeel Manaf, Endalkachew Ashenafi Alemu, Cathrine Broberg Vågbø, Leonardo Martín, Mads Lerdrup, Klaus Hansen, Sinnakaruppan Mathavan, Cecilia Winata, Robert B. Darnell, Peter Aleström, Arne Klungland

## Abstract

Early vertebrate embryogenesis is characterized by extensive post-transcriptional regulation during the maternal-to-zygotic transition. The *N*6-methyladenosine (m^6^A) modifications on mRNA have been shown to affect both translation and stability of transcripts. Here we investigate the m^6^A topology during early vertebrate embryogenesis and its association with polyadenylated mRNA levels. The majority (>70%) of maternal transcripts harbor m^6^A, and there is a substantial increase of m^6^A in the polyadenylated mRNA fraction between 0 and 2 hours post fertilization. Notably, we find strong associations between m^6^A, cytoplasmic polyadenylation and translational efficiency prior to zygotic genome activation (ZGA). Interestingly, the relationship between m^6^A and translation is strongest for peaks located in the 3’UTR, but not overlapping stop codons. Sequence analyses revealed enrichment of motifs for RNA binding proteins involved in translational regulation and RNA degradation. After ZGA, m^6^A seem to diminish the effect of miR-430 mediated degradation. The reported results improve our understanding of the combinatorial code behind post-transcriptional mRNA regulation during embryonic reprogramming and early differentiation.

## Introduction

Post-transcriptional chemical modifications of RNA alter their fate and are important for RNA function. The *N*6-methyladenosine (m^6^A) modification is the most abundant internal mRNA modification, and each mRNA contains on average three to five m^6^A modifications [1, 2]. The fate and life-time of m^6^A containing mRNAs is partly regulated through specific interaction with the YTH-domain containing proteins. RNAs bound by YTHDF2 is transported to decay sites resulting in increased degradation rates [3], while mRNAs bound to YTHDF1 are more efficiently translated through interaction with the translation machinery [4]. In addition to these direct effects, m^6^A can change the secondary structure of transcripts and allow proteins to bind otherwise hidden sequence motifs [5, 6].

Zebrafish is a very attractive system to study embryogenesis and associated processes such as reprogramming and differentiation, including their associated transcriptomic and epigenetic blueprints. Although the zebrafish share embryonic features with other species, embryogenesis is remarkably fast compared with mammals (Fig 1a). The egg is activated upon water contact, and after fertilization and the zygote period (1-cell; 0 to 0.75 hours post fertilization, hpf), the embryo start to divide rapidly during cleavage stages (0.75 to 2 hpf), reaching 64-cells before entering the blastula period (2.25 to 5.25 hpf). During this time span, zygotic genome activation (ZGA, ~3 hpf), occurs. At 4hpf (“sphere”) there is robust expression of many genes important for further developmental progress [7, 8]. The blastula period cells can form embryonic stem cells in culture, similar to the inner cell mass of mouse embryos [9, 10]. Thus, from fertilization until blastula, the germ cells have been reprogrammed to become pluripotent. During the gastrula period (5.25 to 10 hpf), the germ layers are formed. The gastrula period is characterized by more extensive changes in cell division synchrony and cell migration patterns, leading to the formation of the “shield” at 6 hpf.

**Figure 1.**
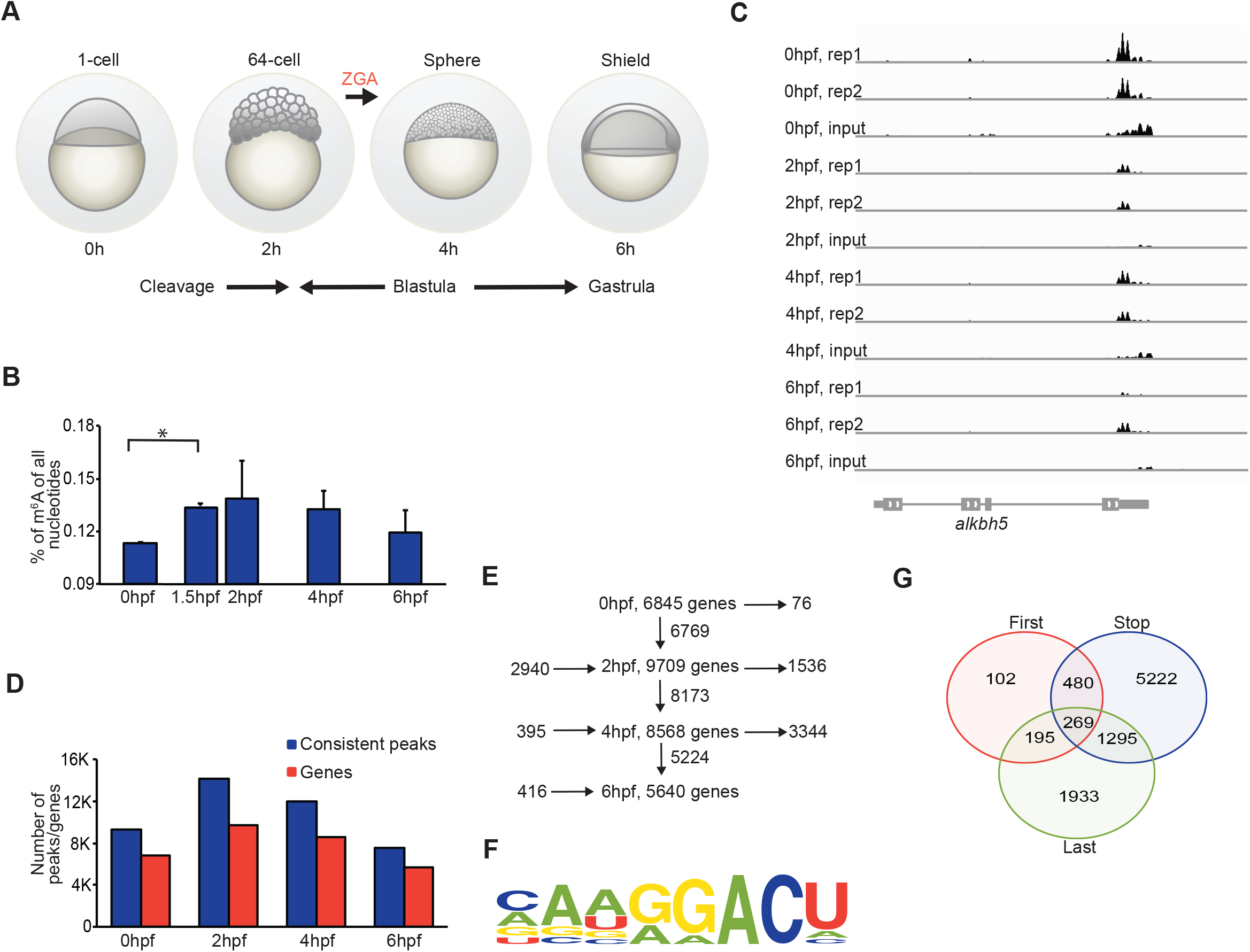
Zebrafish embryogenesis and m^6^A dynamics. A) Four stages of zebrafish early embryogenesis with descriptive names on top, and hours post fertilization (hpf) below embryos. ZGA; Zygotic genome activation. B) Amount of m^6^A in the mRNA fraction during development measured by LC-MS/MS. The bars represent the average of three replicates and error bars depicts one standard deviation. Stages are from left to right; 0 hpf, 1.5 hpf, 2 hpf, 4 hpf and 6 hpf. C) Integrative genome browser (IGV) snapshot showing all m^6^A-RIP-seq replicates, and one selected input sample from each stage. Height of the barplot indicates enrichment level. D) Number of peaks and genes with m^6^A identified by m^6^A-IP-sequencing. E) m^6^A fate map during early embryogenesis. Number of genes that gain m^6^A on the left side, genes with m^6^A in boxes, genes with maintained enrichment below boxes and number of genes with lost m^6^A on the right side. F) Enriched motif found in peak regions in the 0 hpf sample (p-value <0.001, 53% vs 30%, hypergeometric test). G) Venn diagram showing which genes have peaks in the different regions.

Due to the lack of transcription, early embryonic development relies on maternally provided transcripts and proteins [8]. Post-transcriptional mechanisms such as elongation of polyA tails through cytoplasmic polyadenylation (CPA), and mRNA degradation are crucial for normal developmental progression [11]. CPA causes transitioning of maternal transcript from a dormant to an actively translatable state [7, 12, 13]. This mechanism regulates thousands of maternal transcripts [7, 14], including important factors for reprogramming and ZGA, such as *pou5f3* and *nanog* [12, 15]. How these transcripts are targeted for CPA after fertilization remains largely unknown, since the amount of variation explained by *cis* elements is low, and most studies are performed on maturing oocytes [16, 17].

Degradation of maternal RNAs and proteins is achieved both through maternal and zygotic programs [18]. Motifs in the 3’UTR such as AU-rich elements (AREs) together with stabilizing (e.g. HuR/Elavl1) or destabilizing (e.g. Auf1) RNA binding proteins (RBPs) are implicated in regulating the stability of transcripts [19]. After ZGA, degradation of many maternal RNAs is mediated by the microRNA-430 (miR-430) family [20]. However, miR-430 miRNAs have many more predicted targets than those affected by miRNA knock-down [20]. In the cases of both mRNA activation and degradation it therefore remains a possibility that there are additional regulatory layers that determines which transcripts are targeted for degradation, or CPA and translation.

m^6^A is dynamically regulated in the heat shock response and during sporulation in yeast [21, 22]. Recent studies have also indicated a role for m^6^A in regulating pluripotency related genes and the reprogramming process [23-25]. More recently it was reported that m^6^A reader protein Ythdf2 mediates degradation of maternal transcripts in zebrafish embryos [26]. However, other potential effects of m^6^A, such as cytoplasmic polyadenylation and translational efficiency, were not examined.

We therefore found it timely to investigate the role of m^6^A during early embryogenesis in a broader perspective. We performed m^6^A-RNA-immunoprecipitation followed by sequencing (m^6^A-IP-seq), and compared it to multiple published datasets available for the studied developmental period (0 to 6 hpf). We find that m^6^A is a highly prevalent modification on maternally produced RNA transcripts, with an estimate of 75% of the genes being modified at some level, much higher than recent estimates [26]. There is a substantial increase of m^6^A in the mRNA fraction between 0 and 2 hpf. After extensive CPA, m^6^A modified transcripts are translated at a substantially higher level at 2 hpf (64-cell stage) than transcripts without m^6^A, and generally become degraded between 4 and 6hpf (blastula to gastrula stage). Furthermore, between 4 and 6 hpf, m^6^A containing transcripts targeted by miR-430 are more stable than miR-430 targets without m^6^A. These observations warrant further functional studies, and implicate m^6^A in several key molecular processes important for embryonic development, including ZGA and reprogramming.

## Materials and methods

### Zebrafish husbandry and stage selection

Embryos from the AB wild type strain were obtained from the NMBU zebrafish facility where the zebrafish were kept at 28±1°C on a 14-10 hour light-dark cycle at a density of 5-10 fish/L. System water (SW) was prepared from particle and active charcoal filtrated tap water, deionized by reverse osmosis (RO) and kept sterile by UV irradiation. The water was conditioned to a conductivity of 500µS/cm, general hardness (GH) 4-5 and pH 7.5 by addition of 155g synthetic sea salt (Instant Ocean, Blacksburg, USA), 53g sodium carbonate and 15g calcium chloride (Sigma-Aldrich) per liter RO water. Adult fish were fed with Gemma Micro 300 (Skretting, Stavanger, Norway) dry feed twice a day and live artemia (Scanbur, Karlslunde, Denmark) once a day. Health monitoring was by daily inspection, use of sentinel fish sent for pathology (ZIRC, Eugene, Oregon) and water microbiology analysis (NMBU Vetbio, Oslo) every six month. Adult fish were allowed to mate for 30 minutes in standard 1L breeding tanks (Aquatic Habitats, Apopka, FL). Harvested embryos were kept in autoclaved SW at 28 °C, harvested by snap freezing (liquid nitrogen) and visually controlled for stage and lack of abnormalities at the selected time points. The zebrafish facility and the used SOPs, including protocol for fish line maintenance, have AAALAC accreditation (No. 1036), and is approved by the National Animal Research Authority. All experiments were performed according to Norwegian Animal Welfare Act (2009), the EU Directive 2010/63.

### Quantification of global m^6^A by LC-MS/MS

We collected three batches of 50 embryos and isolated total RNA from each using the RNeasy MicroKit (QIAGEN, #74004), in which a needle and syringe and a QIAshredder (QIAGEN, #79654) was used to disrupt the embryos. PolyA^+^ RNA was extracted using Dynabeads^®^ mRNA Purification Kit (Ambion, #61006).

RNA was hydrolyzed to nucleosides by 20 U benzonase (Santa Cruz Biotech), 0.2 U nuclease P1, and 0.1 U alkaline phosphatase (Sigma) in ammonium acetate pH 6.0 and 1 mM magnesium chloride at 37ºC for 40 min, added 3 volumes of methanol and centrifuged (16,000 g, 30 min, 4 ^o^C). The supernatants were dried and dissolved in 50 µl water for LC-MS/MS analysis of m^6^A and unmodified nucleosides. Chromatographic separation was performed on a Shimadzu Prominence HPLC system with a Zorbax SB-C18 2.1×150 mm i.d. (3.5 µm) column equipped with an Ascentis Express C18 150 × 2.1 mm ID (2.7 μm) column equipped with an Ascentis Express C18 5 × 2.1 mm ID (2.7 µm) guard column (Sigma-Aldrich). The mobile phase consisted of water and methanol (both added 0.1 % formic acid), for m^6^A starting with a 5-min gradient of 5-60 % methanol, followed by 5 min re-equilibration with 5 % methanol, and for unmodified nucleosides maintained isocratically with 30 % methanol. Mass spectrometric detection was performed using an MDS Sciex API5000 triple quadrupole (AB Sciex) operating in positive electrospray ionization mode, monitoring the mass transitions 282.1/150.1 (m^6^A), 268.1/136.1 (A), 244.1/112.1 (C), 284.1/152.1 (G), and 245.1/113.1 (U).

### m^6^A-RNA immunoprecipitation and sequencing

We isolated total RNA from four zebrafish developmental stages (1-cell/20 minutes post fertilization, 2, 4 and 6 hpf), in batches of 200 embryos, using TRIzol (Invitrogen, cat.no. 15596-018). We added ERCC spike-in RNA (TermoFisher scientific, cat.no. 4456740) to the trizol. RNA yield and quality can be found in table S1. For each sequencing library two batches of total RNA (from 200 embryos) were merged and enriched for polyA^+^ RNA using Dynabeads^®^ mRNA Purification Kit (Ambion, #61006). Two biological replicates (with 2 × 200 embryos in each) for each stage were used in the m^6^A-IP experiment. The m^6^A-IP experiment was carried out as previously described [27]. Briefly, polyA^+^ enriched RNA was partially fragmented by alkaline hydrolysis, ethanol precipitated and SDS-PAGE size selected for 20-80nt. Part of the fragmented RNA was used as input, to account for changes in polyadenylated mRNA levels, and the rest was immunoprecipitated at 4°C for 2 h using Dynabeads Protein A (Life Technologies, # 10008D)-conjugated anti-m^6^A antibody (Synaptic systems, # 202003). After stringent washing, the bound RNA was eluted with 0.5mg/ml N6-methyladenosine sodium salt (Sigma-Aldrich, # M2780), ethanol precipitated, and resuspended with RNase-free water. The eluted RNA and input RNA were subjected to 3’pre-adenylated DNA linker ligation with T4 RNA ligase2, truncated KQ (NEB, #M0373L), at 16 ^o^C over night. The sequencing library was constructed with bromodeoxyuridine (BrdU)-CLIP protocol described in [28], with improved RT primers as indicated in [27], without crosslinking. The samples were sequenced in four lanes on an Illumina HiSeq 2000. The data is deposited under GEO accession number GSE89815.

### Analyses of m^6^A-IP-seq

The sequencing output from each of the 4 lanes in the sequencing run was trimmed 7 nucleotides in the 5’end, and de-multiplexed (4 samples in each lane). Reads from both m^6^A-IP and input samples were mapped to Zv10 using the STAR aligner [29] with options --seedSearchStartLmax 15 --clip3pNbases 10 --clip5pNbases 10 --outFilterMultimapNmax 20 --outFilterMismatchNoverLmax 0.05 --outFilterMatchNminOverLread 0.0 --outFilterMatchNmin 15 --outFilterScoreMinOverLread 0.0.

Approximately 99% of reads mapped (Table S2). For the input control samples, the number of input reads per ensemble83 gene was obtained using featureCounts v.1.5.0 [30], using default options. The data was normalized as previously described [31] after obtaining normalization factors using calcNormFactors in the edgeR package [32]. For visualization in IGV we made tdf files using IGVTools [33], with options -w 10 --minMapQuality 5, thus only visualizing high confidence reads.

We performed peak calling and detection of differentially methylated genes using ExomePeak [34] (FDR cut-off = 0.05). For all analyses we used the “consistent peaks” dataset (peaks detected in both samples). We selected genes with a 2-fold up or down-regulation for functional annotation analyses.

To generate a complete set of peaks, comparable between samples, we merged peaks found at the different stages (overlapping peaks merged using mergeBed). We then counted the number of high confidence IP and input reads in each peak, and after adjustment for sequencing depth (dividing counts by million high confidence mapped reads in the sample), we calculated the m^6^A-IP/input ratio (adding a value of 0.5 to both the IP and input value to avoid division by zero, and high, but irrelevant ratios for low values). This dataset was used as an estimate of the fraction of transcripts with and without m^6^A (for each gene), and to measure the correlation between samples (using Spearman correlation).

Gene annotations were downloaded from Ensembl83 and we extracted first and last exons, and stop codons from the GTF file using a custom script. Sequences were either obtained directly from Biomart (e.g. 3’UTRs), or extracted using getFasta in the bedTools suite (e.g. m^6^A regions). To construct the m^6^A fate map we intersected bed files from the peak calling using bedTools (intersectBed).

We performed *de novo* motif discovery using HOMER (v. 4.8) [35], motif occurrences using FIMO [36] and discriminant analysis using DREME [37]. For the FIMO analysis we selected a thresholds that for all motifs led to no mismatches from the IUAPAC motif. An exception was made for the U_10_ motif where one mismatch was allowed. EaSeq [38] was used to create heatmaps of m^6^A, stop codons and SAPAS reads. R (https://cran.r-project.org) was used extensively in data mining, statistical testing and for visualization.

### Analyses of published datasets

Datasets were downloaded from public repositories and reads mapped using STAR (v. 2.5.0a). FeatureCounts (v.1.5.0) were used to count reads per gene with parameters according to library type and sequencing technology (see table S3).

## Results

### Global m^6^A levels during embryogenesis are dynamic

We measured the amounts of m^6^A in total and polyA enriched RNA from different early embryonic stages of Zebrafish development using liquid chromatography - mass spectrometry (LC-MS/MS). There was a significant increase of m^6^A (per total unmodified nucleotides) in the polyadenylated RNA fraction between 0 hpf and 1.5 hpf (p = 0.0016 after correction for multiple tests (4 pairwise tests), two-sided t-test, n = 3, 22% increase) (Fig 1b), but there was no change in the amount of m^6^A in total RNA (Fig S1).

### m^6^A-seq and the embryonic m^6^A transcriptome

To gain insight into which polyadenylated mRNAs carry the m^6^A modification we performed m^6^A-RNA-immunoprecipitation followed by high throughput sequencing (m^6^A-IP-seq) in duplicates (400 embryos per replicate), of four developmental time points; 0, 2, 4 and 6 hpf (Fig 1a and see methods and table S1 and S2 for RNA yield and sequencing results). To account for fluctuations in polyadenylated RNA levels, which are substantial during these stages, we also included input controls for each replicate. Thus, in total, the dataset consist of 16 sequencing libraries. Transcript isoform quantification is still imprecise [39], therefore we opted for a gene centric analysis. We refer to “genes” in the following, yet, we are aware that transcripts are the functional unit.

The data were of high quality as evidenced by the similarity between replicates at the single gene level (Fig 1c), and genome wide correlations of peak scores (Fig S2a). Consistent with the mass spectrometry results we found a substantial increase in the number of peaks and genes with m^6^A between 0 and 2 hpf, and a substantial decline between 4 and 6 hpf (Fig 1d). This dynamic was also evident when testing for differentially methylated genes between developmental stages (Fig S2b). We constructed an m^6^A peak diagram of genes found to have m^6^A at one or more of the four developmental stages (figure 1e). This diagram demonstrated that 99% of the 6845 m^6^A containing genes at 0 hpf are also present with m^6^A at 2 hpf, plus an additional 2,940 genes. Subsequently there are very few novel m^6^A containing genes, but a substantial loss between 2 and 6 hpf (*n* = 4,880). Interestingly, there is an 85% overlap between genes that gain m^6^A between 0 and 2 hpf, and those losing m^6^A between 2 and 6 hpf. Thus, a subset of genes displays a highly dynamic pattern, whereas a core group of genes has m^6^A enrichment at all stages.

In several species it has been shown that m^6^A occurs in a RRACU sequence context. We did a *de novo* motif discovery and for all samples (0, 2, 4 and 6 hpf) there were an overwhelming enrichment for motifs that included the RRACU motif (Fig 1f) (e.g. p-value at 0 hpf = e^−480^, binomial distribution). Direct search for the RRACU motif found that this was present in 90.3-91.2% of the peaks. This validates the m^6^A-IP-seq results further, and shows that the preference for the RRACU motif is conserved also in zebrafish.

### Maternal and zygotic transcript display differences in m^6^A levels and frequency

The fraction of maternal genes (defined as all genes expressed at 2 hpf) detected with m^6^A was dependent on the gene expression level cut off used, ranging from 57% at 10 counts per million (CPM, based on input sample gene counts), to 69% at 50 CPM cut off. In contrast, for zygotic genes (expression >10 CPM at 6hpf and >2 log2 fold-change increase between 4 and 6 hpf), only 34% (297 of 873 genes) of the genes were methylated (odd ratio 2.56, 95% CI 2.21-2.97, p-value <2.2e^−16^, Fisher exact test).

It has been shown that the percentage of m^6^A modified transcripts in a particular transcript species (e.g. the number of m^6^A modified and unmodified *nanog*) vary between transcripts [40, 41]. As a rough assessment of this fraction we calculated the m^6^A-IP/input ratio (see methods). We found that the median log2 m^6^A-IP/input ratio for maternal transcripts was 1.33, while it was 1.70 for zygotic transcripts (p-value <2.2e^−16^, Wilcoxon rank sum test). Maternal transcripts are thus more commonly methylated than zygotic, but a smaller percentage of the transcripts (for a particular transcript species) are methylated.

### Genes with dynamic m^6^A status are regulators of embryonic events

We used Metascape to investigate the functionality of genes marked with m^6^A at different stages [42]. We focused on the two most dynamic periods, the increase pre-ZGA and the changes occurring after ZGA by selecting genes that had a m^6^A log2 fold change >2. Consistent with the substantial number of genes that both gain methylation pre-ZGA and lose it post-ZGA, the two groups (up 0-2 hpf and down 4-6 hpf) were enriched for similar terms related to developmental processes (e.g. embryonic morphogenesis and cell fate commitment) (Fig S3a and b). Genes with lower m^6^A level at 6 hpf (relative to 4hpf) were in addition enriched for genes with hydrolase activity and kinase regulators (Fig S3b). In contrast, genes that gain m^6^A after ZGA are most frequently involved in chromatin restructuring and retinol metabolism (Fig S3c). This enrichment fits well with the transition of modified histones and nucleosomes around this time [43, 44], and the necessity of retinoic acid signaling during gastrulation [45].

### m^6^A are preferentially found over or in the vicinity of stop codons

In order to identify the location of m^6^A within genes we intersected peak coordinates with annotated features (>1 bp overlap with 3’UTRs, first and last exons, and stop codons). Because annotations can be incomplete and may not perfectly represent early developmental stages, we used only last exons that had both a 3’UTR and a stop codon associated with them. The results showed that the vast majority of peaks are found in last exons (81-86%), and a minor fraction in first exons (6.2-8.7%), consistent with published data [46]. Between 44 and 49% of the peaks overlapped directly with a stop codon. A gene centered Venn diagram of the position of peaks revealed that most genes with a last exon peak also had a peak overlapping the stop codon (Fig 1g). Similarly, first exon peaks were rarely present without other m^6^A peaks (<10% of first exon peaks). Of note, the antibody used for m^6^A-IP-seq (see methods) does not discriminate between m^6^A and N^6^,2’-O-dimethyladenosine (m^6^Am). The latter modification was recently shown to enhance mRNA stability [47].

Interestingly, we found that the m^6^A/input ratio for peaks stratified by their position in the transcript were distinct in terms of signal strength. Peaks in last exons that did not overlap with a stop codon, had a much lower m^6^A /input ratio than peaks overlapping a stop codon (log2 enrichment of 1.33 and 1.78 at 0 hpf, *n* = 2,512 and 5,469, p-value <2.2e^−16^, Wilcoxon test).

In order to improve the resolution of the spatial relationship between m^6^A, stop codons and polyadenylation sites we visualized last exons with m^6^A and their annotated stop codons using EaSeq [38]. As multiple polyadenylation sites are frequent during embryogenesis [48, 49], we included SAPAS data (sequencing of alternative polyadenylation sites) from corresponding stages to visualize the utilized polyadenylation site [49]. The stop codon was frequently found in the beginning of the last exon (Fig 2a), and overlapped with the normalized m^6^A signal, but not the SAPAS signal, both at 0 and 6 hpf. To investigate if the enrichment at the 5’ end was related to the end of the exon, or the stop codons, we visualized the start and end of each last exon, which again showed high degree of co-occurrence between stop codons and m^6^A, and not with the SAPAS signals (Fig 2b). The results shows that when m^6^A is present at last exons containing a stop codon, it occurs in close proximity to the stop codon, while the distance to the cleavage site vary depending on the length of the last exon.

**Figure 2.**
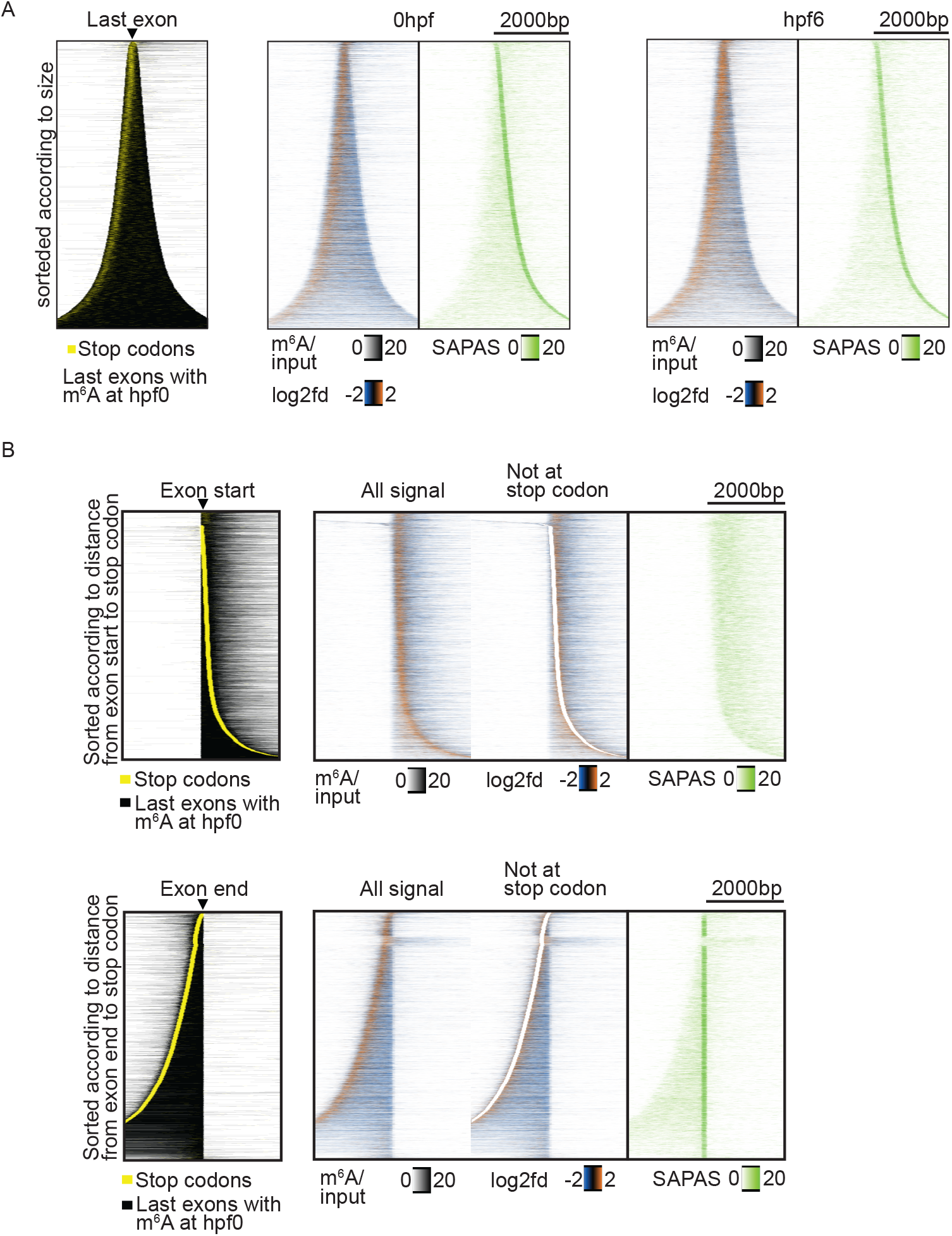
m^6^A enrichments are associated with stop codons. A) Heatmaps of last exons with an annotated stop codon and m^6^A enrichment at 0 hpf. Leftmost heatmap shows the length of the exon in black and the stop codon in yellow. The two next heatmaps display m^6^A enrichment (first) and the density of SAPAS reads (second) at 0 hpf. The same heatmaps are shown for 6hpf. B) Start (upper panel) and end (lower panel) of last exons with an annotated stop codon and m^6^A enrichment at 0 hpf. Leftmost heatmap visualizes the stop codon position, the next three m^6^A enrichment, m^6^A outside the stop codon and SAPAS read density (green).

### Comparison with total RNA m^6^A IP sequencing

In a recent publication ribosome depleted RNA was used to IP m^6^A during zebrafish embryogenesis (in contrast to our polyA+ selected RNA) [26] (GSE79213). To evaluate if our result were in any way biased by the large fluctuations in polyadenylated RNA levels during development we wanted to compare our results from polyA+ selected RNA with that of rRNA depleted RNA. To test if we “missed out” on a lot of genes because of polyA+ RNA selection we identified a cohort of 877 genes that had very high total RNA levels compared to polyA+ levels (at 0 hpf), suggesting they might not be captured during polyA+ selection. Even though underrepresented, 315 of these were still found to be methylated in our dataset (at 2 hpf), and 562 were found to be unmethylated. Only 10 of these unmethylated (10 out of 562) were detected in the total m^6^A CLIP-seq experiment, and these genes included ribosomal RNA (5.8S), ribozymes (Nuclear RNAase P) and Metazoa Signal Recognition Particle RNA. Thus, non-coding RNA components of the ribosome. Three genes were protein coding according to annotation. From these analyses we conclude that;

i. Most genes underrepresented in polyA+ sequencing are not methylated (562 of 877 genes).
ii. Most of those that are methylated are still detected as such (315/325).
iii. Only a handful of underrepresented genes are detected by total RNA based methods, and the majority of these are ribosomal components not translated (but still found in RPF-seq because they are components of the ribosome).

### m^6^A, cytoplasmic polyadenylation and translational efficiency

To gain insight into the role of m^6^A in RNA biology during early vertebrate embryogenesis we took advantage of the rich repertoire of different sequencing experiments for zebrafish embryonic stages that are publicly available (see table S3 for references and accession number).

### PolyA-tail length and mRNA levels

The pre-ZGA stages of development are characterized by extensive polyA tail dynamics, in particular the addition of adenosines to the already existing polyA tails of maternal transcripts, a process referred to as cytoplasmic polyadenylation (CPA) [16]. To analyze a possible relationship between m^6^A and polyA tail length we first used direct measurement of the polyA tail length obtained from poly(A)-tail length profiling by sequencing (PAL-seq) experiment [13]. We found only a small difference between m^6^A and non-m^6^A transcripts both at 2 hpf (24.5 and 23.2 nt, respectively), and at 6hpf (39.7 and 36.9 nt, respectively). However, there is substantial CPA prior or just after fertilization [7, 50], before the first time point in the PAL-seq study (2 hpf). Therefore, it remained a possibility that m^6^A can regulate CPA either in the oocyte, or immediately after fertilization. As an indirect measure of CPA we used mRNA-seq levels from published mRNA-seq datasets [7, 51]. Since transcripts here are selected through hybridization between the polyA tail and oligo(dT) beads, and only low levels of transcription occur, the bulk of mRNA abundance increase must be due to CPA [8, 31].

Due to large difference in average polyadenylated mRNA levels between m^6^A and non-m^6^A genes, we stratified the genes into comparable subgroups with similar levels. For every subgroup, m^6^A genes increase substantially more than non-m^6^A genes pre-ZGA (Fig 3a). To assess at which time point the increase occur we used our previously published dataset, which included several early time points [7]. We found a robust difference between the 1-and 16-cell stage (median log2 fold-change 1.51 and 0.88, n = 7148 and 5970, expression >10 reads at 1-cell stage, p < 2.2e^−16^, Wilcoxon rank sum), but only a minor difference between 16-and 128-cell (median log2 fold-change 0.25 and 0.24, n = 7148 and 5970, expression >10 reads at 1-cell stage). Next, we extracted genes with relatively higher methylation level at 2 hpf compared to 0 hpf, and found that these had a significantly higher increase pre-ZGA compared to genes with m^6^A at 0 hpf (1.55 and 1.95 log2 fold-change, p = 2.7e^−11^, n = 5,143 and 737, Kolmogorov–Smirnov test, only genes expressed >50 CPM at the 1-cell stage).

**Figure 3.**
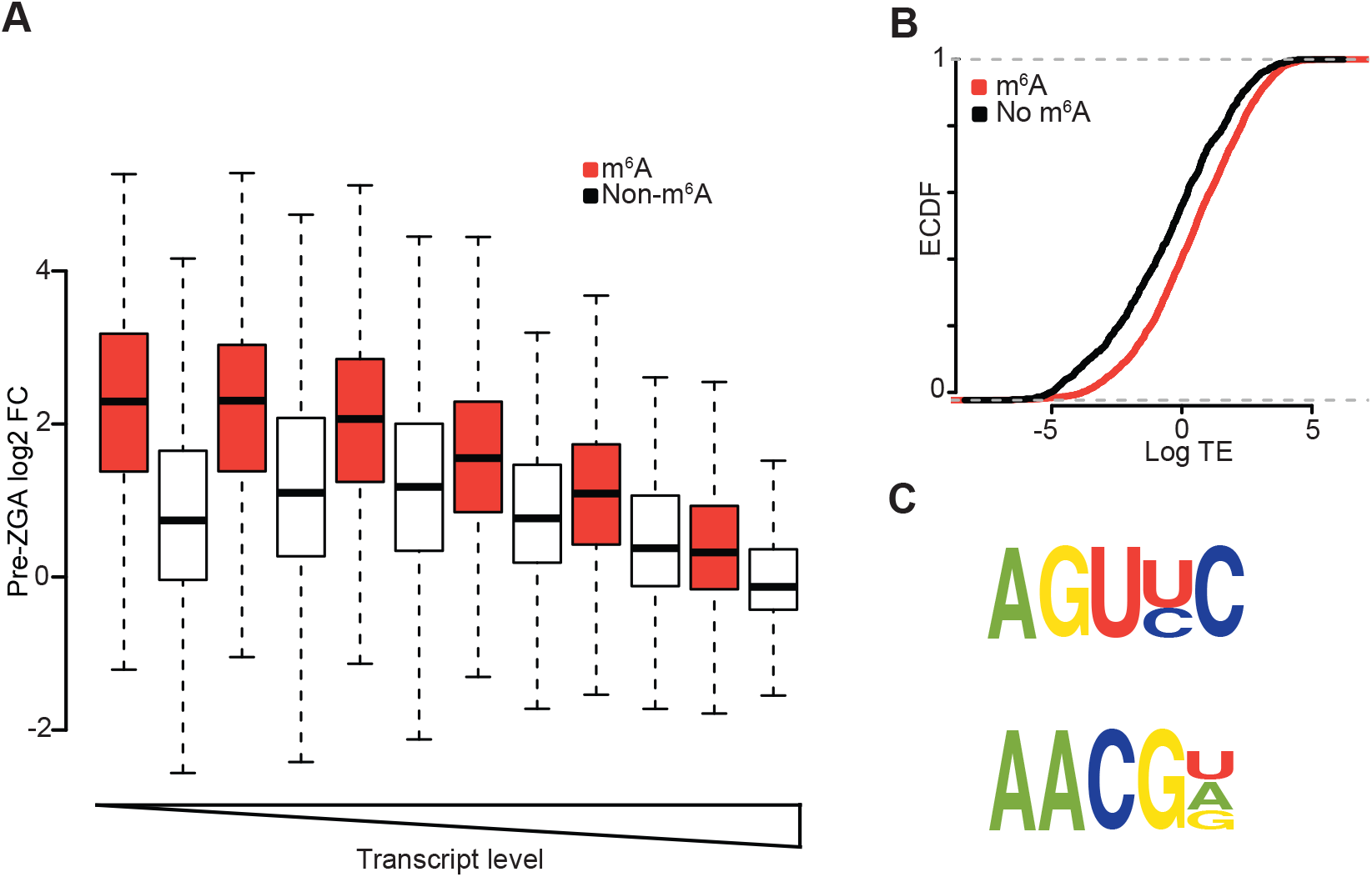
m^6^A and RNA stability, cytoplasmic polyadenylation and translation. A) Boxplot of genes in different expression classes, from low to high polyA+ mRNA levels (left to right), divided by m^6^A status. B) ECDF plot of TE values for genes with and without m^6^A at 2 hpf. C) Motif discovery results from discriminant analysis of 3’UTRs using DREME.

### m^6^A is associated with higher translational efficiency pre-ZGA

There is a strong relationship between polyA tail length and translational efficiency during embryogenesis [13]. However, m^6^A has also been shown to enhance translation through the m^6^A binding protein YTHDF1 [4]. To evaluate a possible relation between m^6^A and translation during early embryogenesis we generated a dataset with genes that had translational efficiency (TE) values at three stages (2, 4 and 6 hpf, *n* = 3,541), using published ribosome protected fragments sequencing (RPF-seq) data [13]. Through intersections with our m^6^A peak data we found a large difference in the TE between m^6^A (*n* = 2,648) and non-m^6^A genes (*n* = 893) at 2 hpf (median TE 0.55 and −0.31, p-value <2.2e-^16^, Kolmogorov–Smirnov test) (Fig 3b). At 6 hpf, the translation efficiency for m^6^A and non-m^6^A was more similar (0.91 and 0.54 for m^6^A and non-m^6^A, respectively). Also of interest, we found that genes with m^6^A located in the last exon, but not overlapping a stop codon, had much higher TE than genes with m^6^A over the stop codon or in the 5’UTR (TE 0.85, 0.52 and 0.32 for last, stop codon and first exon peaks respectively).

### m^6^A is associated with sequence motifs bound by translational regulators

As shown earlier the RRACU motif was the most overrepresented motif within peak regions. To gain a deeper insight to the m^6^A sequence context we performed a more extensive sequence analysis. We performed discriminant analysis between the 3’UTR of m^6^A and non-m^6^A genes, to potentially detect novel motifs that are different between the two groups. The two top motifs enriched in m^6^A genes at 2 hpf were AGU(U/C)C (p-value = 3.4e^−156^, 81% vs 65%), and AACG(U/A) (p-value = 3.0e^−121^, 74% vs 58%) (Fig 3c). The former motif bear close resembles to a reported Dazl binding motif [52], while the latter is bound by Unr, with highest affinity when present together with an upstream purine rich motif [53]. Both Dazl and Unr have been shown to be involved in translational regulation. Dazl promote translation, probably through elongation of the polyA tail [52, 54], while Unr interact with CCR4 and PABP in translation dependent decay [55].

### mRNA degradation and translational inhibition through miR-430 is modulated by m^6^A

During the first 2 hours of embryo development in zebrafish there are only trace levels of transcription [8], which make this time frame ideal to study the stability of maternally produced transcripts. Using two complementing total RNA-seq datasets [12, 56], we found no significant differences in the pre-ZGA stability of m^6^A and non-m^6^A genes at 2 hpf (median log2 fold-change - 0.21 and −0.28) [56].

After ZGA, miRNAs of the miR-430 family clear the maternal transcriptome [18, 20]. We found that miR-430 targets are enriched for m^6^A at 2 hpf (78% and 67% of genes with and without the miR-430 seed had m^6^A, respectively, p <2.2e^−16^, OR 1.82, 95% CI 1.65-2.01, Fisher exact test). To investigate if m^6^A might modify the outcome of miR-430 seed presence in target mRNAs, we examined the changes in total RNA levels [12]. Notably, there was a significant difference between miR-430 target genes, with and without m^6^A, towards enhanced stability if they contained an m^6^A residue at 2 hpf (median log2 fold-change −0.94 and −1.80, n = 822 and 575, p <0.001, Kolmogorov-Smirnov test) (Fig 4a). Interestingly, for m^6^A and non-m^6^A genes without the miR-430 seed had smaller differences (median log2 fold-change −0.34 and −0.60).

**Figure 4.**
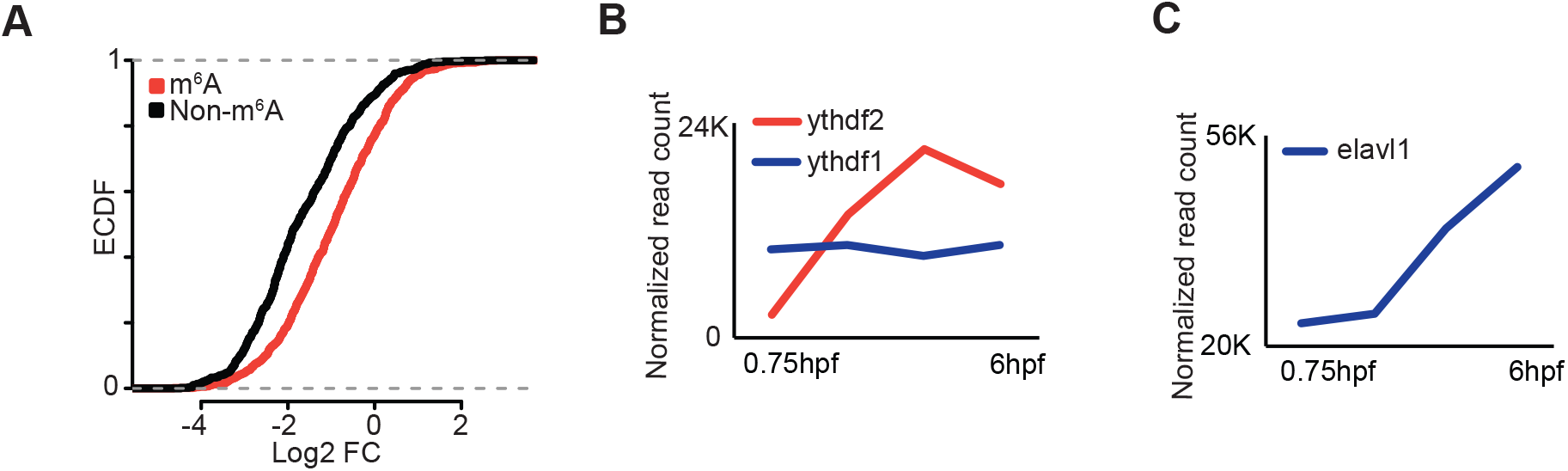
m^6^A modulates mRNA fate of miR-430 targeted transcripts. A) ECDF plot of log2 fold-change values post-ZGA from total RNA-sequencing for genes containing the miR-430 seed and stratified by presence or absence of m^6^A enrichment. B) and C) Expression of different m^6^A readers during development.

### Expression of writers, readers and erasers of m^6^A during development

Proteins involved in m^6^A functionality can be divided into writers, readers and erasers. Importantly, all known genes involved in m^6^A regulation are conserved in zebrafish. Interestingly, both the writer *mettl14* and the eraser *alkbh5* have robust increase pre-ZGA (Figs S4a and b). This suggests a highly dynamic m^6^A state, with frequent methylation and demethylation. Notably, we observed high levels of the m^6^A reader and translational regulator *ythdf1* at all stages (Fig 4b). In contrast, *ythdf2* increase from relatively low levels pre-ZGA and reaches its top level at 4.3hpf (Fig 4b). Another gene suggested to be involved in m^6^A gene regulation, *elavl1*, has robust increase in expression from ZGA and onwards (Fig 4c) [3, 57].

## Discussion

Our study improves the understanding of two key processes during embryogenesis, post-transcriptional regulation of maternal transcripts, and miR-430 mediated regulation post-ZGA. There are, however, important questions that remain unanswered. First, do m^6^A affect CPA only, and thus indirectly translation, or both CPA and translation independently? The earlier timing of CPA, and the delay in effect on translation, but not CPA, suggests that CPA is the primary driver. An exception from this might be for temporarily restricted peaks located away from stop codons, but still in last exons. They were most strongly associated with translation, yet they display lower IP/input ratio, suggesting that they are very potent signals for translational increase. Preliminary analysis using multiple linear regression found that m^6^A presence were an significant independent predictor, together with polyA tail length, in predicting TE, however the main proportion of TE variation is explained by polyA tail length.

Second, it needs to be proven functionally which proteins are involved in the regulation of m^6^A levels and effects on mRNA dynamics. It seems likely that *ythdf1*, already expressed at high levels at fertilization and previously shown to enhance translation [4], is involved. Further, *ythdf2* and *elavl1* can alter the stability of transcripts [3, 58], and their expression profiles suggest a role during or after ZGA. Indeed, it was recently shown that *ythdf2* knock-out cause developmental delay after ZGA, probably caused by reduced degradation of maternal transcripts and lower expression of zygotic transcripts [26]. However, only a small number of high confidence *ythdf2* targets were identified (n = 135). We could not detect any general effects on total RNA stability prior to ZGA, consistent with the expression pattern of *ythdf2* (Fig 4f). However, genes with m^6^A at 2 hpf were more strongly degraded post-ZGA, in line with the observed effect of *ythdf2* binding. Yet, degradation is not a general effect of m^6^A, since genes that retained their m^6^A modification at 6 hpf were more stable than non-m^6^A genes post-ZGA. Our results are consistent with a more complex model of m^6^A function, in which the mRNA fate depends on the combinatorial effect of multiple RNA binding proteins, directly to m^6^A and/or due to effects on secondary structure. The outcome of m^6^A is therefore likely to vary in time and space. Our observations indicate that prior to ZGA the predominant effect of m^6^A is enhancement of translation, and after ZGA, on stability.

Another gene possibly associated with post-ZGA RNA regulation is Elavl1. Although pulled down in an m^6^A bait approach and experimentally linked to m^6^A [3, 57], it has been questioned if Elavl1 binds m^6^A directly [59]. However, structural imprints *in vivo* have shown that m^6^A alter the secondary structure of mRNAs [6], and that the altered structure can affect the ability of RNA binding proteins to target their sequence motifs [5]. This might be the case for the observed association between m^6^A and Elavl1. Similarly, we found several sequence motifs for protein factors such as translational regulators Dazl and Unr, enriched in the vicinity of m^6^A (Fig 3c). Enhanced RNA binding of these proteins might also contribute to the RNA dynamics we observed. It is likely that the relative position of m^6^A, protein binding sequences and secondary structures predict the mRNA fate.

Future work will also be needed to determine and understand the onset of high m^6^A levels in early embryogenesis. The relatively lower levels of m^6^A in MII oocytes (3820 genes) reported in *Xenopus* [60], indicates that the increase in m^6^A occurs later than the MII stage. Further studies using recently developed methods to determine differences in structure and co-occurrence with RNA binding proteins will increase the understanding of how m^6^A contributes to RNA biology. The current work lays an interesting fundament for future mechanistic studies and shed light into the combinatorial code of mRNA regulation during early vertebrate embryogenesis.

## Acknowledgements

The LC-MS/MS analyses were provided by the Proteomics and Metabolomics Core Facility (PROMEC), Norwegian University of Science and Technology (NTNU). PROMEC is funded by the Faculty of Medicine at NTNU and Central Norway Regional Health Authority. EAA is supported by Funding from the European Union Seventh Framework Programme (FP7-PEOPLE-2013-COFUND) under grant agreement n° 609020 - Scientia Fellows. We acknowledge the support from the Norwegian Cancer Society, the Research Council of Norway and the Research Program of the EEA/Norway Grants (grant number POL/NOR/196258/2013). This work was performed on the Abel Cluster, owned by the University of Oslo and the Norwegian metacenter for High Performance Computing (NOTUR), and operated by the Department for Research Computing at USIT, the University of Oslo IT-department. http://www.hpc.uio.no/

## Authors’ contributions

HA and AK designed the study. HA performed all analyses except for EaSeq analyses, and wrote the manuscript. CBV performed LC-MS/MS. EAA was responsible for m^6^A-IP-sequencing. ML performed analyses using EaSeq. DE, LM and AM performed embryo collection and RNA isolation procedures. CW performed morpholino experiments and contributed polysome sequencing data. SM contributed with polysome sequencing data. PA contributed with zebrafish facilities and RD m^6^A-IP-seq facilities. AK supervised the project. All authors contributed to discussions and approved the final manuscript.

## Data availability

The data is deposited under GEO accession number GSE89815. All data points and datasets used in this study are found in supplementary file 1.

