## Supplementary material for "*N*6-methyladenosine dynamics during early vertebrate embryogenesis"

Figure S1

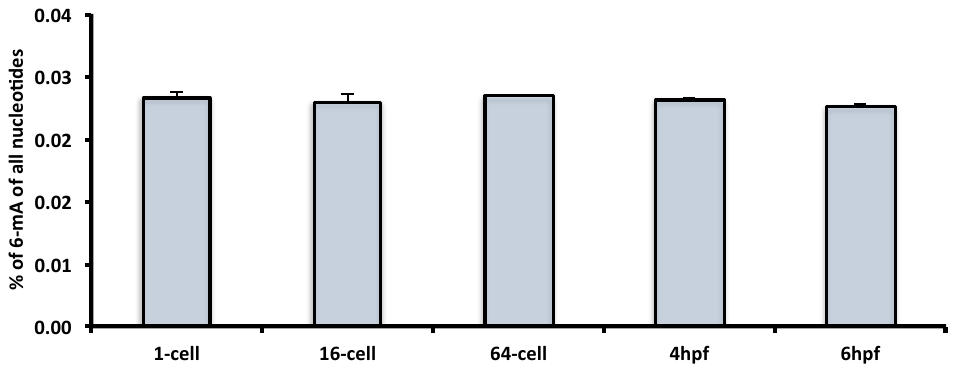

Figure S2a

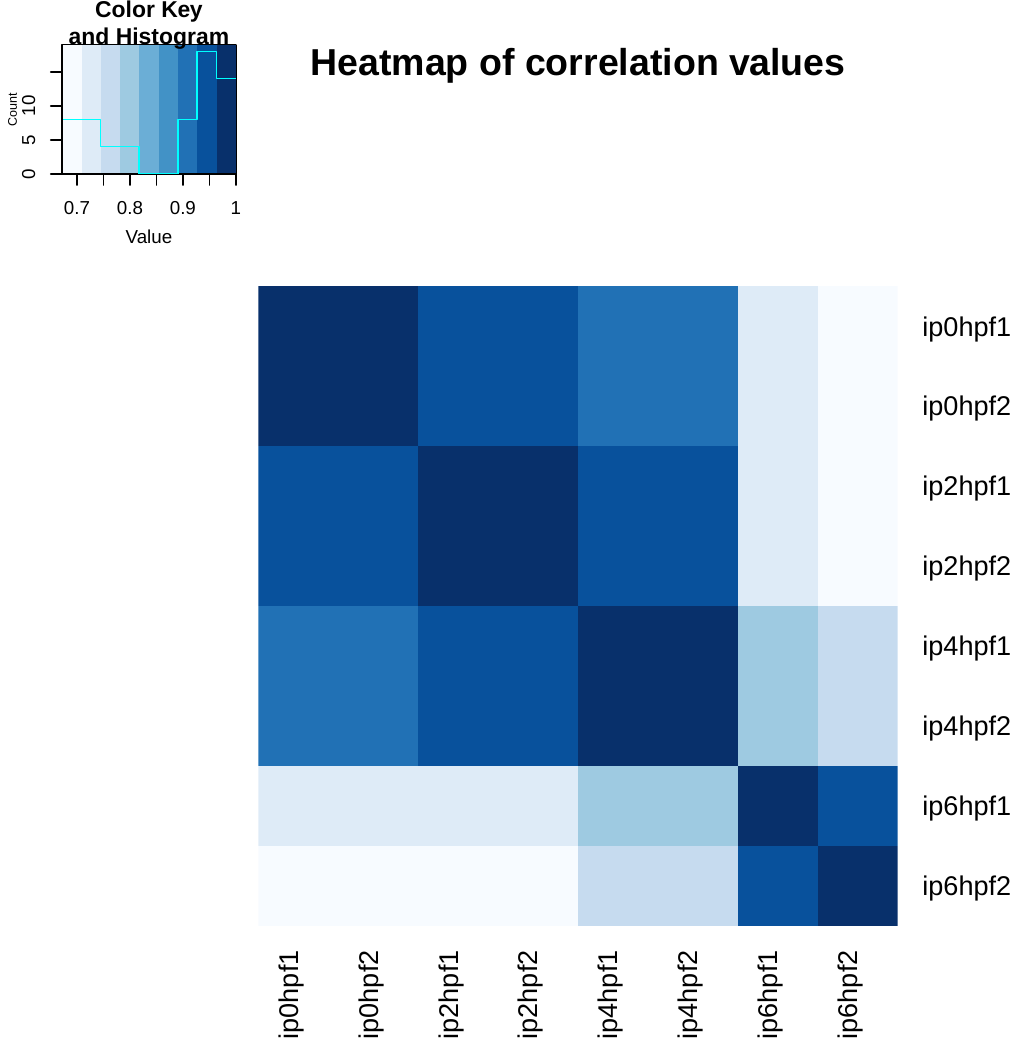

Figure S2b

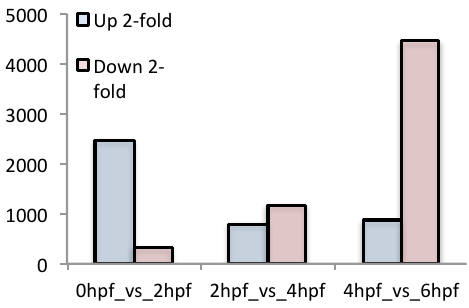

Figure S3a

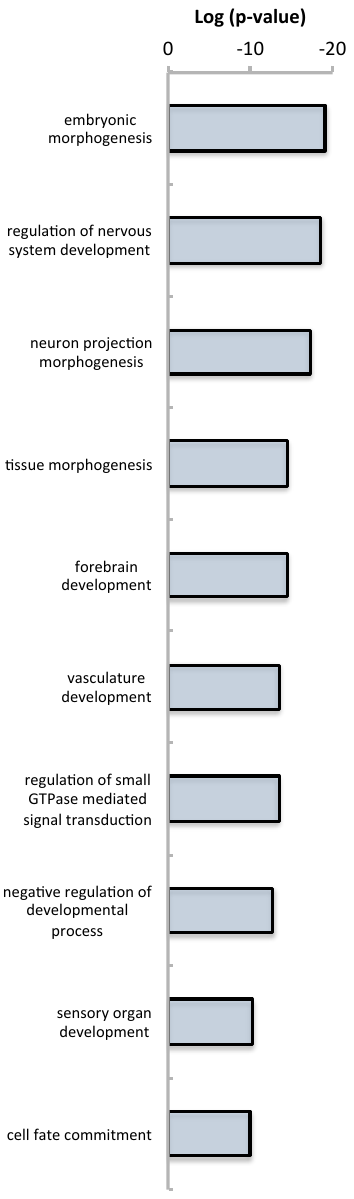

Figure S3b

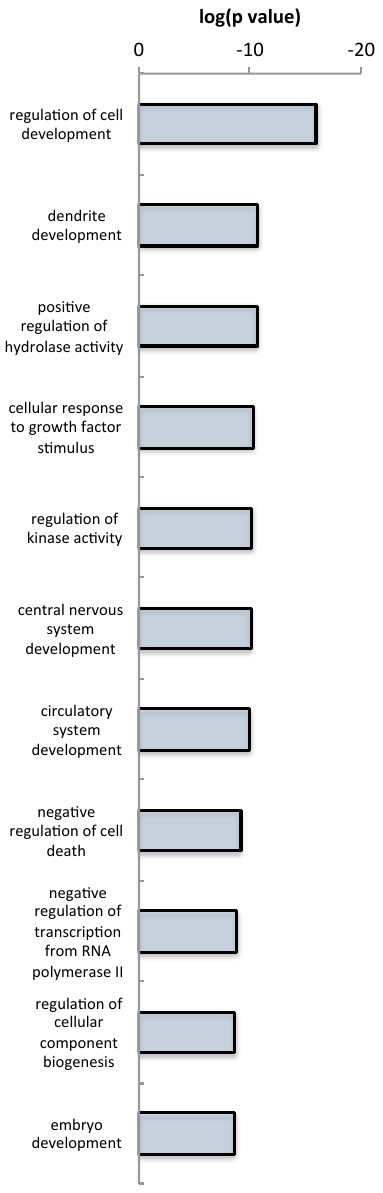

Figure S3c

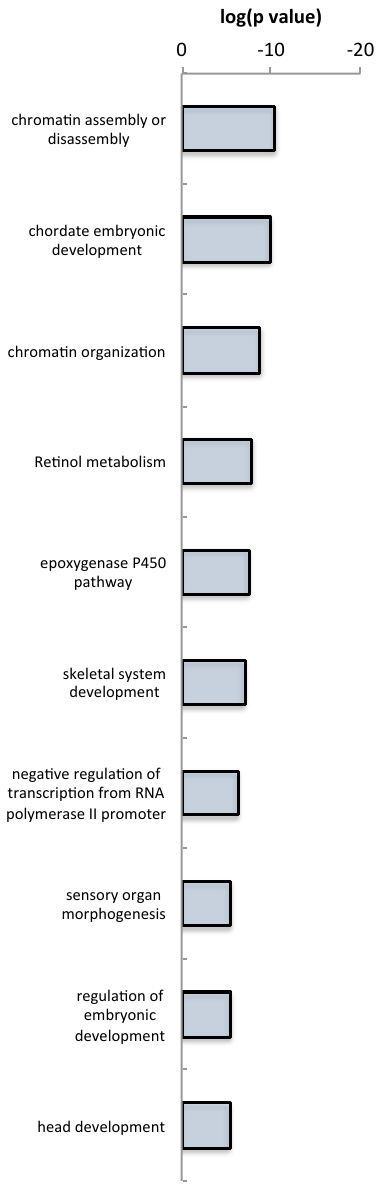

Figure S4A

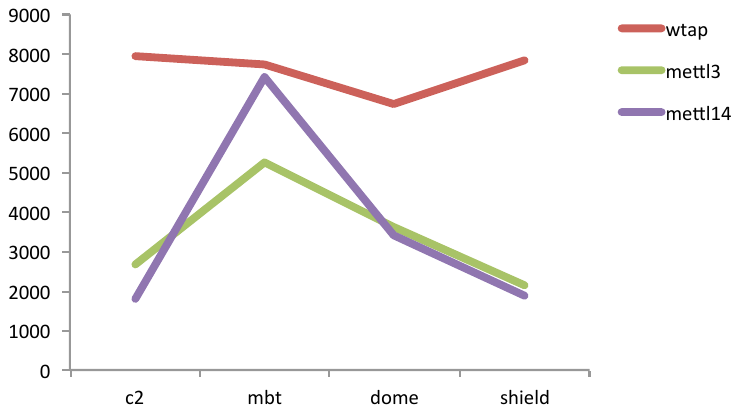

Figure S4B

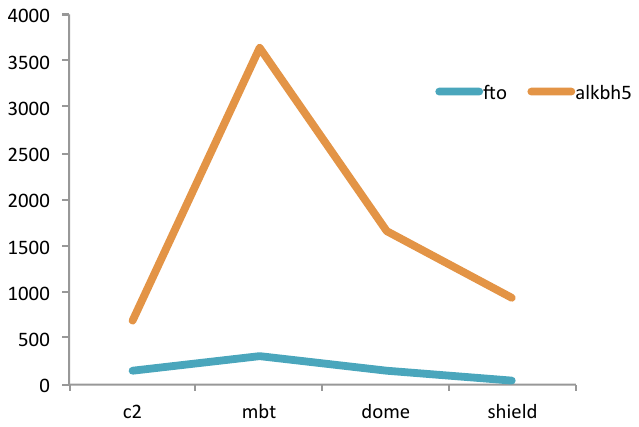

Table S1. RNA yield

| Sample | Nanodrop Conc ng/µl | Total amount in 50 ul | ug | ng Per embryo estimate |
| --- | --- | --- | --- | --- |
| Zv_1-Cell (1st Rep)_total-RNA_18.8.15 | 595 | 29747 | 29,7 | 148,7 |
| Zv_1-Cell (2nd Rep)_total-RNA_18.8.15 | 511 | 25558 | 25,6 | 127,8 |
| Zv_1-Cell (3rd Rep)_total-RNA_18.8.15 | 693 | 34658 | 34,7 | 173,3 |
| Zv_1-Cell (4th Rep)_total-RNA_18.8.15 | 753 | 37672 | 37,7 | 188,4 |
| Zv_64-Cell (1st Rep)_total-RNA_18.8.15 | 334 | 16694 | 16,7 | 83,5 |
| Zv_64-Cell (2nd Rep)_total-RNA_18.8.15 | 691 | 34539 | 34,5 | 172,7 |
| Zv_64-Cell (3rd Rep)_total-RNA_18.8.15 | 768 | 38395 | 38,4 | 192,0 |
| Zv_64-Cell (4th Rep)_total-RNA_18.8.15 | 824 | 41194 | 41,2 | 206,0 |
| Zv_Sphere (1st Rep)_total-RNA_18.8.15 | 748 | 37392 | 37,4 | 187,0 |
| Zv_Sphere (2nd Rep)_total-RNA_18.8.15 | 679 | 33956 | 34,0 | 169,8 |
| Zv_Sphere (3rd Rep)_total-RNA_18.8.15 | 843 | 42159 | 42,2 | 210,8 |
| Zv_Sphere (4th Rep)_total-RNA_18.8.15 | 718 | 35881 | 35,9 | 179,4 |
| Zv_Shield (1st Rep)_total-RNA_18.8.15 | 825 | 41274 | 41,3 | 206,4 |
| Zv_Shield (2nd Rep)_total-RNA_18.8.15 | 729 | 36441 | 36,4 | 182,2 |
| Zv_Shield (3rd Rep)_total-RNA_18.8.15 | 775 | 38727 | 38,7 | 193,6 |
| Zv_Shield (4th Rep)_total-RNA_18.8.15 | 815 | 40771 | 40,8 | 203,9 |

Table S2. Generated and mapped reads

| Sample | Reads | Mapped,Unique | Mapped, Multiple | Total, Mapped | %Mapped | %Unique |
| --- | --- | --- | --- | --- | --- | --- |
| Input_0hpf_1 | 38 833 209 | 30 330 872 | 8 377 538 | 38 708 410 | 99,7 | 78,1 |
| Input_0hpf_2 | 35 954 915 | 27 862 595 | 7 994 297 | 35 856 892 | 99,7 | 77,5 |
| Input_2hpf_1 | 37 647 859 | 28 902 656 | 8 639 229 | 37 541 885 | 99,7 | 76,8 |
| Input_2hpf_2 | 38 284 919 | 30 021 498 | 8 157 381 | 38 178 879 | 99,7 | 78,4 |
| Input_4hpf_1 | 40 154 421 | 29 700 914 | 10 237 760 | 39 938 674 | 99,5 | 74,0 |
| Input_4hpf_2 | 39 200 596 | 29 463 421 | 9 514 111 | 38 977 532 | 99,4 | 75,2 |
| Input_6hpf_1 | 40 062 481 | 29 774 592 | 9 887 104 | 39 661 696 | 99,0 | 74,3 |
| Input_6hpf_2 | 69 785 511 | 53 272 879 | 15 856 738 | 69 129 617 | 99,1 | 76,3 |
| IP_0hpf_1 | 38 406 033 | 22 016 142 | 15 983 163 | 37 999 305 | 98,9 | 57,3 |
| IP_0hpf_2 | 37 079 070 | 25 234 079 | 11 591 426 | 36 825 505 | 99,3 | 68,1 |
| IP_2hpf_1 | 41 617 061 | 29 379 822 | 11 925 247 | 41 305 069 | 99,3 | 70,6 |
| IP_2hpf_2 | 37 560 404 | 27 807 920 | 9 491 664 | 37 299 584 | 99,3 | 74,0 |
| IP_4hpf_1 | 44 530 500 | 28 539 075 | 15 557 738 | 44 096 813 | 99,0 | 64,1 |
| IP_4hpf_2 | 41 770 496 | 25 977 815 | 15 289 557 | 41 267 372 | 98,8 | 62,2 |
| IP_6hpf_1 | 43 698 133 | 22 106 010 | 20 861 388 | 42 967 398 | 98,3 | 50,6 |
| IP_6hpf_2 | 41 019 829 | 23 231 493 | 17 148 755 | 40 380 248 | 98,4 | 56,6 |

Table S3. Sequencing datasets and processing steps

| **Type** | **First author** | **Year** | **Stages** | **Processing** | **GEO, SRA or ENA accession #** |
| --- | --- | --- | --- | --- | --- |
| Total RNA-seq | Lee | 2013 | 2, 4 and 6 hpf | STAR --outFilterMultimapNmax 10 --outFilterMismatchNoverLmax 0.05 --seedSearchStartLmax 15 --clip3pNbases 10 --clip5pNbases 10 --outFilterMatchNminOverLread 0.0 --outFilterMatchNmin 15 --outFilterScoreMinOverLread 0.0 | GSE47558 |
| Total RNA-seq | Vesterlund | 2011 | 1-cell, 16-cell, 512-cell, epiboly50 | Mapped using Tophat (v.2.0.13) and Bowtie (v. 1.1.2.0) with default options –C, --quals and –G. | ERP000635 |
| PAL-seq | Subtelny | 2014 | 2, 4 and 6 hpf | Downloaded processed data from GEO | GSE52809 |
| mRNA-seq | Pauli | 2012 | 2/4-cell, 3hpf, Sphere, shield | Reads mapped with STAR, parameters; --outFilterMultimapNmax 20 --outFilterMismatchNoverLmax 0.05. Reads were counted with featureCounts with parameters -s 2, and -p. | GSE32898 |
| mRNA-seq | Aanes | 2011 | 1-cell, 16-cell, 128-cell, 3.5 hpf, 5.3 hpf | Used previously processed data (Aanes et al., 2013) | GSE22830 |
| SAPAS | Li | 2012 | 0, 4 and 6 hpf | STAR options; --seedSearchStartLmax 15 --clip3pNbases 10, --clip5pNbases 10, --outFilterMultimapNmax 20,  --outFilterMismatchNoverLmax 0.05,  --outFilterMatchNmin 15. | SRA036536 |
