## Supplementary material for "*N*6-methyladenosine dynamics during early vertebrate embryogenesis": Legends supplementary material

**Legends, supplementary figures and tables**

**Figure S1. m^6^A in total RNA**. m^6^A per 1000 nucleotides in total RNA during development measured by LC-MS/MS.

**Figure S2. m^6^A-RIP-seq reproducibility and change**. A) Heatmap of correlation values between samples. B) Number of differentially methylated genes between different developmental stages.

**Figure S3. Functional annotation of m^6^A genes.** Functional annotation results from Metascape of genes with higher methylation levels 2 hpf (to 0 hpf) (A), lower methylation levels at 6hpf (to 4 hpf) (B) and higher methylation levels at 6 hpf (relative to 4 hpf) (C). Top 10 enriched terms are shown.

**Figure S4. Expression of m^6^A writers and erasers.** Line plot showing the expression of genes involved in methylation and demethylation of transcripts. A) Expression of m6A writers wtap, mettl3 and mettl14. B) Expression m^6^A erasers fto and alkbh5.

**Legends, supplementary tables**

Table S1. RNA yield. Amount of RNA retrieved from each batch of 200 embryos.

Table S2. Generated and mapped reads from sequencing.

Table S3. Dataset and their associated processing steps used in this article.
